# Historical data reveals earlier flowering under environmental change

**DOI:** 10.64898/2025.12.16.694749

**Authors:** Elizabeth Stunz, Ante Stattin Gustavsson, Christine D. Bacon

## Abstract

**Premise:** Global temperatures have risen ∼1.1°C since the 1800s and the mean annual air temperature in the Arctic has increased nearly four times the global average. Temperature has a significant effect on fundamental life-cycle events of flora and fauna. With temperatures increasing, Arctic plants are expected to shift their active periods accordingly, with a shorter flowering period as later flowering plants converge with flowering timing of early flowering plants.

**Methods:** Nearly 120 years of herbarium records and collection data were leveraged to identify flowering shifts in the Arctic-alpine plant *Oxyria digyna* using climatic data and linear regressions.

**Results:** We found that Alpine, Boreal Arctic, and Arctic Tundra populations differed in their flowering time response to increased air temperature. The Boreal Arctic population showed a significant advancement in flowering time as related to mean summer and June temperature increases, while the flowering time of Arctic Tundra and Alpine populations had a shift in flowering timing in response to June temperature change.

**Conclusions:** Our results identify biodiversity response to climate change and determine which populations have greater adaptive potential to respond to rapidly changing abiotic conditions in the Arctic and beyond.

## Introduction

The global Earth surface temperature has risen ∼1.1°C since the mid-to-late 1800s (Lee et al., 2023). While climate is changing on a global scale, the greatest changes in air temperature and its yearly variation have occurred in the Arctic (Lee et al. 2023). For example, average air temperature in the Arctic has increased at nearly four times the global average (Rantanen et al., 2022). As the Arctic warms, thinner snowpacks, increased permafrost thaw, and accelerated melting and retreat of glaciers, ice sheets, and sea ice exacerbate environmental change in the region (Lee et al., 2023). Contemporary environmental change in the Arctic has altered the physiology and phenology, the core timing of seasonal patterns, of many species. For instance, the termination of hibernation of male and female Arctic ground squirrels in Alaska have gradually diverged due to warmer winter temperatures (Chmura et al., 2023), and juvenile Arctic charr have experienced increased growth rates linked to increased autumn temperatures (Kotowych et al., 2023). Shifts in the green-up date of many Arctic plants have also been documented, such as the sedge *Carex aquatilis* and the grass *Arctophila fulva*, which advanced by 16 days between 1945 – 2016 and showed a strong correlation with number of thaw degree days each year (days above 0°C; Andresen et al., 2018).

The phenology of plant life cycle events, such as green-up and flowering time, is crucial to maximize reproductive success (González-Suárez et al., 2020). Most genes that affect flowering timing are strongly regulated by abiotic conditions, such as day length and temperature (Capovilla et al., 2015). A changing climate has led to detectable shifts in the timing of these events in some plant species (Walther et al. 2002), and responses vary across and within regions (Love & Mazer, 2021). For example, flowering phenology of eight out of ten alpine species was found to be highly temperature sensitive, especially during peak flowering at mid-summer (Hülber et al. 2010). The phenology of Arctic plants at colder, high latitude sites showed a greater sensitivity to temperature than those at warmer, lower latitude sites (Prevéy et al., 2017). Further, phenological sensitivity to shifts in climate (i.e. temperature and precipitation) can differ depending on resource availability and site elevation (Ahmad et al., 2021; Yao et al., 2021). An improved understanding of phenological response at the community-, species-, and population-levels are essential to better understand how species and ecosystems are currently responding and will respond in the future to our changing climate.

To explore climate change response in plants, here we focus on *Oxyria digyna* (Polygonaceae), an herb found in both alpine and Arctic regions. The species is broadly distributed across the northern hemisphere at elevations between 0 – 4500m, making it an ideal species to investigate response to climate change. Genomic analyses infer an origin in alpine regions of eastern Asia, and subsequent migration to northern Asia at ca. 11 Ma., North America at ca. 4 Ma, Europe at ca. 1.96 Ma, and Greenland at ca. 0.91 Ma (Wang et al., 2016). While its geographic distribution is broad the species is habitat specific, occurring only in tundra and on talus slopes, rock outcrops, gravel, and bluffs (Mooney & Billings, 1961). A later-flowering species, *O. digyna* typically flowers from June to August and clinal variation in flowering phenology suggests strong local adaptation (Mooney & Billings, 1961). While Arctic tundra and alpine populations did not initiate flowering significantly earlier in short-term warming experiments (Heide, 2005), a two-decade study showed *O. digyna* flowered earlier under warmer spring temperatures and was significantly correlated to the date of snowmelt (Bjorkman et al., 2015). While these studies have detected phenological responses at shorter temporal scales (Heide, 2005; Mooney & Billings, 1961; Bjorkman et al., 2015), there remains a knowledge gap regarding phenological responses of *O. digyna* over longer temporal scales (>20 years).

As century-scale datasets are rarely available, one approach to identify long-term effects is to leverage natural history collections. For instance, collections from herbaria track flowering through specimen metadata across time (Davis et al., 2015; Lang et al., 2019; Auffret, 2021). By leveraging herbarium collections and occurrence records, we here aim to identify shifts in flowering time of *O. digyna* across its range over ca. 120 years. Based on results at short time scales (Heide, 2005; Mooney & Billings, 1961; Bjorkman et al., 2015), we hypothesize that there are spatial and temporal differences in flowering time of *O. digyna* across the Arctic-alpine latitudinal gradient. Further, we hypothesize that a longer temporal scale will reveal strong response of flowering to temperature increase. From these hypotheses, we expect that flowering time shifts are correlated to rising temperatures, and therefore we predict the largest shifts in flowering timing to occur in the Arctic tundra, as the greatest temperature increases are occurring there. Our statistical regression results identify flowering time shifts and determine their significant correlation to mean summer temperature and June temperature increases from 1901-2021. These results provide unequivocal evidence of biodiversity response to climate change, leveraging long-term data sources that reinforce the importance of natural history collections as fundamental resources for biodiversity science.

## Materials and Methods

We compiled occurrence records of *Oxyria digyna* from the Global Biodiversity Information Facility, a global network of biodiversity data (GBIF, 2024). Only occurrences with geographic coordinates and linked herbarium specimen records were used to determine if the specimen was in flower, so flowering shifts could be precisely tracked over time and space. Data was cleaned for biases (i.e. country centroids and museums) using the CoordinateCleaner R package v.3.0.1 (Zizka et al., 2019). The final dataset comprised of “GBIF occurrence ID”, “year”, “month”, “day”, “flowers present”, “latitude”, “longitude” and “dataset” (the herbarium where the specimen is preserved) of each record.

Elevation data for each occurrence record was inferred from its geographic coordinates using functions from the elevatr R package v.0.99.0 (Hollister et al., 2017) in R 4.3.3. Two Application Programming Interfaces, to allow communication between different software types (Johl, 2022), were called in elevatr to infer elevation data: Elevation Point Query Service (EPQS) and the Amazon Web Services (AWS). EPQS is fast and highly accurate (root mean2 error = 0.53m), but only covers the United States, while AWS has global coverage and 15 different resolution options (Dees, 2017). To account for this, both rasters were processed in elevatr. The difference between the two elevation datasets was calculated (AWS – EPQS = 36,3m) and the AWS dataset was used due to its higher resolution [AWS resolution = 7 (grid size: 611.5m at 60°N].

Following definitions in Arft et al. (1999) and Stunz et al. (2022), *O. digyna* specimens were categorized into one of three populations (alpine, Boreal Arctic, and Arctic Tundra). Occurrences below 60°N were categorized as Alpine, occurrences at 60°N - 68°N as Boreal Arctic, and occurrences above 68°N as Arctic Tundra, as treeline frequently occurs at or near 68°N. Occurrences above 700 meters between 60°N - 68°N were also categorized as Arctic Tundra.

Temperature data was obtained from the Climatic Research Unit CY4.06 dataset (Harris et al., 2022), which contains gridded monthly temperature logs from 1901 to 2021. The data was reformatted from netCDF4 into CSV using the ncdf4 R package v.1.22 (Pierce, 2023). The geosphere R package v.1.5-18 (Hijmans et al., 2022) was used to determine the closest available grid containing a temperature data point. A yearly mean of summer temperatures (June – August) and mean June temperature, respectively, was calculated for each herbarium specimen and attributed to each occurrence record.

Linear regression analyses were conducted using lme4 R package v.1.1-35.3 (Bates et al., 2015) and p<0.05 was considered statistically significant (Di Leo & Sardanelli, 2020). Year, mean June temperature, and mean summer temperature were tested as explanatory variables for the first flowering occurrence in the four datasets (All, Alpine, Boreal Arctic, and Arctic Tundra). Spatial autocorrelation of samples violates an assumption of linear models, so spatial autoregressive (SAR) models were created with the spatialreg R package v.1.3-2 (Bivand et al., 2021) to account for spatial autocorrelation in the data. Neighbourhood groupings for the spatial autoregression were designed to test multiple distance thresholds, spanning from 0 – 2500m, with 250m increments, and flowering occurrences within 250m distance thresholds were grouped. The Pr(>|z|) statistic and Nagelkerke pseudo-R2 value were used to test the p-value and R2 value, respectively, of each model. The model with the best fit was selected based on the lowest Akaike information criterion (AIC) value. For regression plots, SAR model regression lines, based on fitted values, show how well the model predicts the dependent variable response considering spatial relationships between data points.

## Results

After cleaning the occurrence data, a total of 2,243 cleaned GBIF records spanning 119 years were retained for analysis (Figure 1; Table 1). The Arctic Tundra population included 857 records from 1902 to 2020 (except for years 1907, 1909, 1912, 1916, 1918, 1920, 1922, 2014, and 2018 when no specimens adhering to criteria were recorded), the Boreal Arctic population contained 421 occurrences from 1901 to 2019 (excluding years 1902, 1906, 1909, 1911, 1912, 1914, 1915, 1929, 1949, 1980, 1992, 2013, and 2016), and the Alpine population contained 965 occurrences from 1901 to 2020 (excluding years 1902, 1912, and 1917) (Table 1). After filtering occurrences with missing temperature data, the following first flowering occurrences per year were retained for each respective subset for regression analyses: All (n = 330), Arctic Tundra (n = 108), Boreal Arctic (n = 106), and Alpine (n = 116).

**Table 1.**
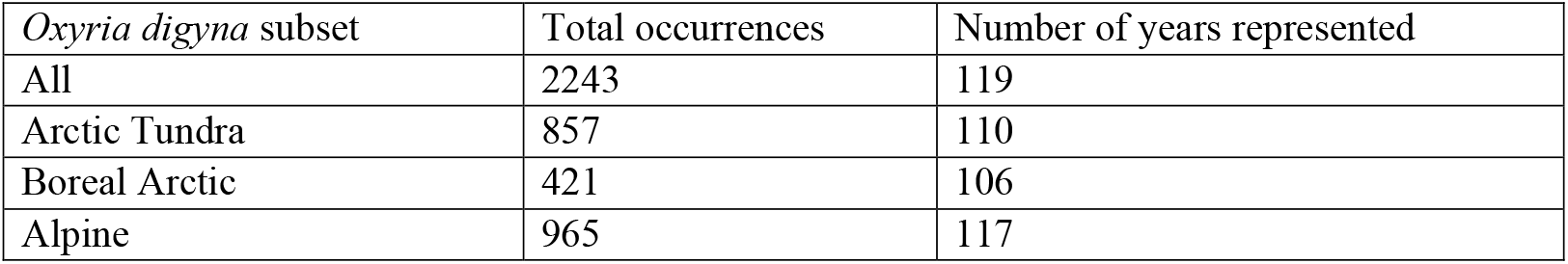
Total occurrences over time for each subset of *Oxyria digyna* occurrences.

| *Oxyria digyna* subset | Total occurrences | Number of years represented |
| --- | --- | --- |
| All | 2243 | 119 |
| Arctic Tundra | 857 | 110 |
| Boreal Arctic | 421 | 106 |
| Alpine | 965 | 117 |

**Figure 1.**
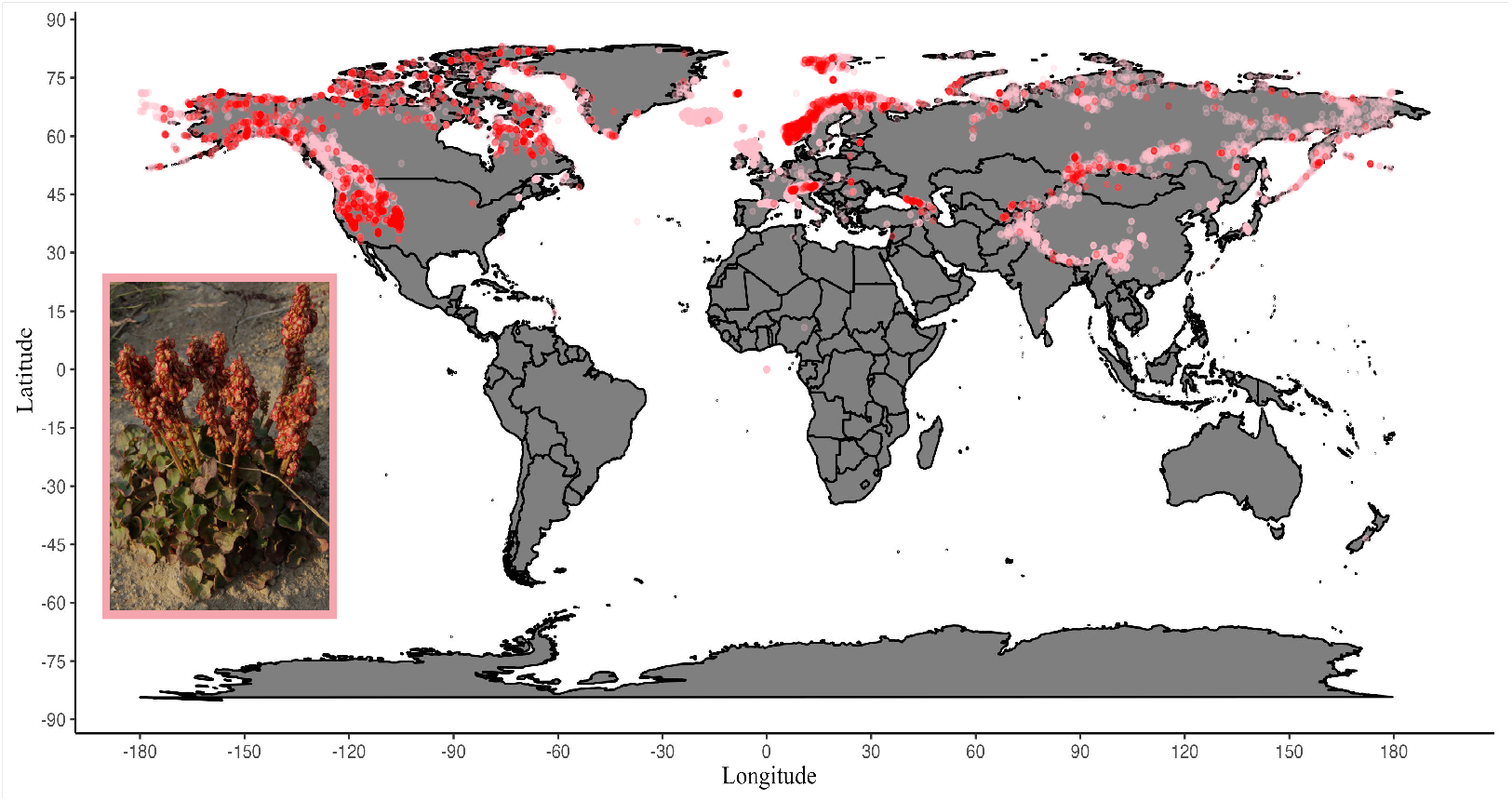
Distribution of *Oxyria digyna* geographic occurrence data. Pink dots indicate all *O. digyna* occurrences available at GBIF and red dots indicate occurrences of records with flowering phenology data. An image of a mature *Oxyria digyna* individual in an Arctic tundra habitat in Danmarkshavn, Greenland is inset in the bottom left. Photo by Elise Biersma.

For all *Oxyria digyna* occurrences, the linear regression model had a strongly significant earlier flowering shift over time (Table 2; effect = -0.093, p = <0.001, R2 = 0.033, AIC = 2771.6). Regressions with mean summer temperature (Table 2; effect = -0.119, p = 0.599, R2 = -0.002, AIC = 2783.2) and mean June temperature (Table 2; effect = -0.311, p = 0.134, R2 = 0.004, AIC = 2781.2) showed a shift to earlier flowering, but it was not significant, and correlations were weak. The spatial autoregressive (SAR) model between year and flowering time of all occurrences showed significantly earlier flowering over time (Table 3; effect = -0.081, p = 0.003, R2 = 0.086, AIC = 2755.7) and in relation to June temperature (Table 3; effect = -0.606, p = 0.026, R2 = 0.075, AIC = 2759.7). Earlier flowering as related to mean summer temperature (Table 3; effect = -0.445, p = 0.140, R2 = 0.068, AIC = 2762.4) was also identified but was not significant.

**Table 2.** Linear regression model statistics for each independent variable (year (1901-2021), mean summer temperature (°C), and mean June temperature (°C) as related to first day of flowering for each *Oxyria digyna* subset. The most supported model (lowest Aikake Information Criterion (AIC) value is in bold).

| *Oxyria digyna* subset | Linear model independent variables | Effect | *p*-value | R<sup>2</sup> | AIC |
| --- | --- | --- | --- | --- | --- |
| All | Year | -0.093 | <0.001*** | 0.033 | 2771.6 |
|  | Mean summer temperature (°C) | -0.119 | 0.599 | -0.002 | 2783.2 |
|  | Mean June temperature (°C) | -0.311 | 0.134 | 0.004 | 2781.2 |
| Arctic Tundra | Year | -0.053 | 0.192 | 0.007 | 874.13 |
|  | Mean summer temperature (°C) | -0.020 | 0.955 | -0.009 | 875.87 |
|  | Mean June temperature (°C) | -0.078 | 0.798 | -0.009 | 875.8 |
| Boreal Arctic | Year | -0.078 | 0.138 | 0.012 | 913.28 |
|  | Mean summer temperature (°C) | -1.842 | 0.004** | 0.068 | 907.04 |
|  | Mean June temperature (°C) | -1.731 | <0.001*** | 0.096 | 903.79 |
| Alpine | Year | -0.134 | 0.003** | 0.066 | 978.76 |
|  | Mean summer temperature (°C) | -0.483 | 0.316 | <0.001 | 986.67 |
|  | Mean June temperature (°C) | -0.812 | 0.074 | 0.019 | 984.44 |

**Table 3.** Spatial Autoregressive (SAR) model statistics for each independent variable (year (1901-2021), mean summer temperature (°C), and mean June temperature (°C) as related to first day of flowering for each *Oxyria digyna* subset. The most supported model (lowest Aikake Information Criterion (AIC) value is in bold).

| *Oxyria digyna* subset | SAR model independent variable | Effect | *p*-value | R<sup>2</sup> | AIC |
| --- | --- | --- | --- | --- | --- |
| All | Year | -0.081 | 0.003** | 0.086 | 2755.7 |
|  | Mean summer temperature (°C) | -0.445 | 0.140 | 0.068 | 2762.4 |
|  | Mean June temperature (°C) | -0.606 | 0.026* | 0.075 | 2759.7 |
| Arctic Tundra | Year | -0.011 | 0.780 | 0.090 | 867.66 |
|  | Mean summer temperature (°C) | 0.022 | 0.962 | 0.090 | 867.73 |
|  | Mean June temperature (°C) | 0.097 | 0.805 | 0.090 | 867.68 |
| Boreal Arctic | Year | -0.077 | 0.125 | 0.065 | 910.41 |
|  | Mean summer temperature (°C) | -1.806 | <0.001*** | 0.135 | 902.1 |
|  | Mean June temperature (°C) | -1.605 | <0.001*** | 0.163 | 898.62 |
| Alpine | Year | -0.101 | 0.018* | 0.177 | 967.13 |
|  | Mean summer temperature (°C) | -0.787 | 0.120 | 0.155 | 970.19 |
|  | Mean June temperature (°C) | -0.942 | 0.037* | 0.169 | 968.29 |

For the Arctic Tundra population, a weakly correlated, non-significant advance in flowering over time was found using the linear regression model (Table 2; effect = -0.053, p = 0.192, R2 = 0.007, AIC = 874.13). Linear regressions also demonstrated non-significant and weakly correlated relationships for earlier flowering as related to mean summer temperature (Table 2; effect = -0.020, p = 0.955, R2 = -0.009, AIC = 875.87) and June temperature (Table 2; effect = -0.078, p = 0.798, R2 = -0.009, AIC = 875.80) in the Arctic Tundra population. The SAR model between year and flowering time showed a non-significant advance in flowering (Table 3; effect = -0.011, p = 0.780, R2 = 0.090, AIC = 867.66), while respective SAR models with mean summer temperature (Table 3; effect = 0.022, p = 0.962, R2 = 0.090, AIC = 867.73) and June temperature (Table 3; effect = 0.097, p = 0.805, R2 = 0.090, AIC = 867.68) variables showed non-significant later flowering.

For the Boreal Arctic population, the linear regression model showed earlier flowering over time, but the shift was not significant (Table 2; effect = -0.078, p = 0.138, R2 = 0.012, AIC = 913.28). Linear regressions for both mean summer temperature (Table 2; effect = -1.842, p = 0.004, R2 = 0.068, AIC = 907.04) and June temperature (Table 2; effect = -1.731, p = <0.001, R2 = 0.096, AIC = 903.79) showed a highly significant earlier flowering shift. Using SAR modelling, a weakly correlated, non-significant shift to earlier flowering was found over time (Table 3; effect = -0.077, p = 0.125, R2 = 0.065, AIC = 910.41), and highly significant relationships were found between mean summer temperature (Table 3; effect = -1.806, p = <0.001, R2 = 0.135, AIC = 902.10) and June temperature (Table 3; effect = -1.605, p = <0.001, R2 = 0.163, AIC = 898.62) and earlier flowering, respectively.

For the Alpine population, significantly earlier flowering over time (Table 2; effect = -0.134, p = 0.003, R2 = 0.066, AIC = 978.76) and non-significant earlier flowering shifts in relation to mean summer temperature (Table 2; effect = -0.483, p = 0.316, R2 = <0.001, AIC = 986.67) and June temperature (Table 2; effect = -0.812, p = 0.074, R2 = 0.019, AIC = 984.44) were found using linear regression models. Spatial autoregressive models showed significantly earlier flowering in relation to year (Table 3; effect = -0.101, p = 0.018, R2 = 0.177, AIC = 967.13) and June temperature (Table 3; effect = -0.942, p = 0.037, R 2 = 0.169, AIC = 968.29), as well as non-significant earlier flowering related to mean summer temperature (Table 3; effect = -0.787, p = 0.120, R2 = 0.155, AIC = 970.19).

## Discussion

Global climate change is causing significant shifts in fundamental life cycle event of flora and fauna. As mean annual air temperature in the Arctic has increased by nearly four times the global average (Rantanen et al., 2022), we expect Arctic biodiversity to be the most severely affected. A convergence of flowering times of earlier-flowering and later-flowering plants in the Canadian Arctic has been identified, as later-flowering species bloom earlier and shift the timing of flowering more than earlier-flowering species (Panchen et al., 2025). Here, we used nearly 120 years of herbarium records and collection data to determine whether temperature increase led to a shift to earlier flowering of *Oxyria digyna* across the Arctic-alpine latitudinal gradient. We expected earlier flowering timing across the gradient, with more rapid shifts in the Arctic Tundra due to amplified temperature increases.

Using both linear and SAR regressions, a significant relationship between earlier flowering in *O. digyna* and an increase in both June temperature and mean summer temperature was observed in the Boreal Arctic (Figure 2). Boreal Arctic individuals flowered significantly earlier (1.7 days per °C increase) in response to increased June temperature over 120 years using the SAR model. Phenology and growth of Arctic plants have been strongly associated with summer temperatures (Bjorkman et al., 2015; Oberbauer et al., 2013; Prevéy et al., 2017), and we show Boreal Arctic individuals are more sensitive to mean summer temperature increases (Table 3). *Oxyria digyna* has varying response to temperature at the individual-level. Short-term studies show no phenological response to temperature (Mooney & Billings, 1961), while Bjorkman et al. (2015) found earlier flowering correlated to higher spring temperatures at decadal scales. Differences in temperature sensitivities of species at warmer and colder sites in the Arctic tundra have been documented, where higher latitude plants at colder sites are more sensitive to temperature (Prevéy et al., 2017). In contrast, we found that Arctic individuals show greater temperature sensitivity at lower latitude sites below treeline as Arctic Tundra *O. digyna* had no significant shift under June or summer temperature increases (Table 3).

**Figure 2.**
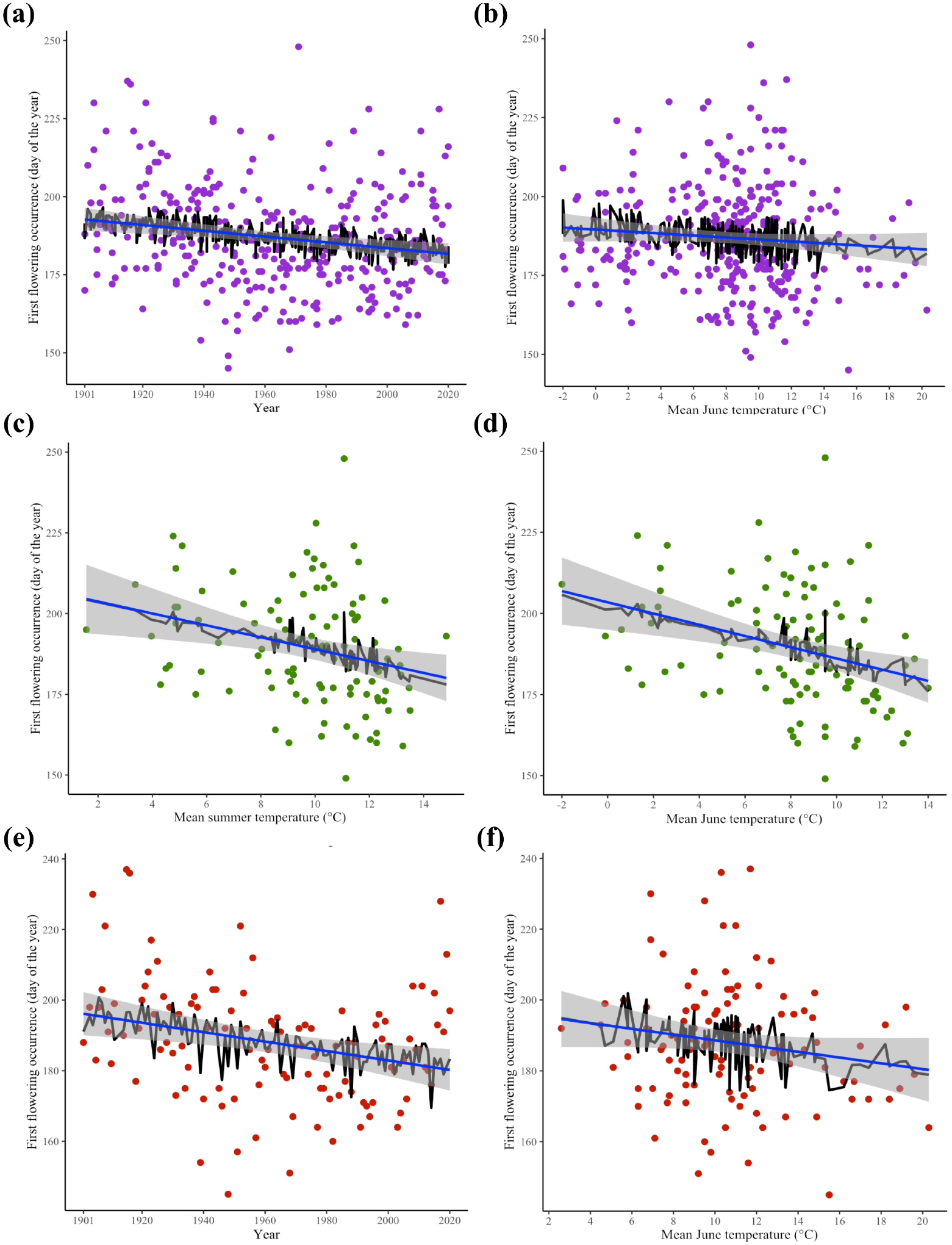
Regression plots of significant spatial autoregressive (SAR) models for the first day of flowering (from 1901-2021) as related to (a) year (1901-2021) and (b) mean June temperature (°C) of all *Oxyria digyna* specimens (indicated by purple dots); (c) mean summer temperature (°C) and (d) mean June temperature of Boreal Arctic *O. digyna* (green dots); and (e) year (1901-2021) and (f) mean June temperature (°C) of Alpine *O. digyna* (red dots). In each respective plot the linear regression line is in blue, and the SAR model regression line based on fitted values is in black.

Alpine *O. digyna* flowering occurred nearly one day earlier per degree Celsius increase in June temperature (Figure 2), indicating a significant shift to earlier flowering. This suggests temperature changes alone will not drive alpine *O. digyna* populations to higher elevations, although the altered population dynamics due to upward shifts of treeline (Dirnböck et al. 2011) and increased species richness of high-elevation habitats linked to warming (Steinbauer et al. 2018) may negatively affect their dispersal and resilience. The Arctic Tundra individuals are found north of treeline, Phenological sensitivity differences while Boreal Arctic individuals occur below treeline. This ecological barrier can restrict gene flow between populations across the barrier, as has been found for other wind-pollinated Arctic plant species (Stunz et al., 2022), and influences population connectivity and adaptive evolution as well.

The difference in temperature sensitivity north and south of treeline supports previous identification of different ecotypes, or locally adapted populations, specific to the alpine and Arctic areas in North America (Mooney & Billings, 1961). Alpine and Arctic *O. digyna* ecotypes differ in morphology and photoperiod requirements for flowering, with shorter day flowering response decreased along a cline as home site latitude increased (Mooney & Billings, 1961; Heide et al., 2005). Phenological sensitivity differences may be due to genomic adaptations formed through a lack of gene flow and/or other evolutionary processes like genetic drift influencing populations north and south of treeline (Rana et al., 2023).

Our results indicate that spatial relationships and other variables are likely influencing flowering timing as well as climate variables, and future research should include more environmental variables to further untangle flowering shifts of populations and/or ecotypes in response to environmental change. While herbarium data may be spatially and temporally biased for various reasons, such as sample site accessibility, herbarium specimens fill gaps in field observation records across climatic and temporal scales to identify phenological response to climate change (Davis et al., 2015).

Here, we identify significant shifts in the flowering timing of *O. digyna* correlated to changes in temperature, with the strongest shift found for Boreal Arctic *O. digyna* in relation to rising June and summer temperatures. With the rapidly changing climate conditions society is currently facing, many ecosystems are at risk of losing genotypes, ecotypes, and species. Understanding differential responses within species to a changing climate is important to determine the potential for short-term evolutionary response and their persistence in warming Arctic and alpine regions.

## Acknowledgements

We thank Søren Faurby and members of the Gothenburg Global Biodiversity Centre for valuable feedback on this work.

## Author Contributions

Elizabeth Stunz: conceptualization, methodology, data analysis; writing – review and editing; Ante Stattin Gustavsson: conceptualization, data curation, data analysis, methodology, writing – review and editing. Christine D. Bacon: conceptualization; funding acquisition; project administration; methodology; writing – review and editing.

## Data Availability Statement

R code and occurrence data can be found at https://github.com/estunz/Oxyria_digyna_phenology

